# Gut Bacteria Encode Reductases that Biotransform Steroid Hormones

**DOI:** 10.1101/2024.10.04.616736

**Authors:** Gabriela Arp, Angela Jiang, Keith Dufault-Thompson, Sophia Levy, Aoshu Zhong, Jyotsna Talreja Wassan, Maggie Grant, Yue Li, Brantley Hall, Xiaofang Jiang

**Affiliations:** Department of Cell Biology and Molecular Genetics, University of Maryland, College Park, College Park, Maryland, USA; National Library of Medicine, National Institutes of Health, Bethesda, Maryland, USA; Division of Molecular and Cellular Biology, Eunice Kennedy Shriver National Institute of Child Health and Human Development, National Institutes of Health, Bethesda, Maryland, USA; Department of Computer Science, Maitreyi College University Of Delhi, Delhi, India; Department of Chemistry and Biochemistry, University of Maryland, College Park, College Park, Maryland, USA; Center for Bioinformatics and Computational Biology, University of Maryland, College Park, College Park, Maryland, USA

**Keywords:** steroid hormones, reductases, metabolism, microbial biotransformation, progesterone

## Abstract

The metabolism of steroids by the gut microbiome affects hormone homeostasis, impacting host development, mental health, and reproductive functions. In this study, we identify the Δ^4^-3-ketosteroid 5β-reductase, 3β-hydroxysteroid dehydrogenase/Δ^5-4^ isomerase, and Δ^6^-3-ketosteroid reductase enzyme families encoded by common human gut bacteria. Through phylogenetic reconstruction and mutagenesis, We show that 5β-reductase and Δ^6^-3-ketosteroid reductase have evolved to specialize in converting diverse 3-keto steroid hormones into their 5β- and Δ^6^-reduced derivatives. We also find that the novel 3β-hydroxysteroid dehydrogenase/Δ^5-4^ isomerase is fused with 5β-reductase in multiple species, streamlining the multi-step conversion of pregnenolone, a steroid hormone precursor, into epipregnanolone. Through metagenomic analysis, we reveal that these enzymes are prevalent in healthy populations, being enriched in females over males. These findings provide the molecular basis for studying microbial steroid metabolism in the gut, offering insights into its potential impact on hormonal health in hosts, especially in the context of women’s health.

## Introduction

Gut microbial metabolism of steroids, such as bile acids, cholesterol, and hormones, is increasingly recognized for its profound impact on human health. Recent studies have explored this through the identification of bacterial enzymes that metabolize steroids in the gut, including the cholesterol dehydrogenase *ismA* ^*1*^, steroid sulfotransferase BtSULT ^2^, bile acid hydrolase BSH ^3^, the *bai* bile acid operon ^4^, and steroid-17,20-desmolase *desAB* ^*5*^. The characterization of these enzymes has deepened our understanding of the microbiome’s role in maintaining steroid homeostasis and its implications for various health conditions, including cardiovascular disease ^6^, pathogen resistance ^7^, prostate cancer ^8^, and colorectal cancer ^9^.

Steroid hormones, including glucocorticoids, mineralocorticoids, androgens, estrogens, and progestogens, are essential for regulating stress responses, immunity, reproductive functions, and metabolic pathways, and maintaining their proper levels is critical for physiological homeostasis ^10–14^. This intricate balance is further influenced by the gut microbiome, which interacts dynamically with these compounds. As part of the natural cycling of these metabolites, hormones are secreted into the gut in bile, where they can impact gut functions, be excreted as waste, or be reabsorbed and circulated back into the body ^15^. Research indicates that significant amounts of steroid hormones, particularly during pregnancy, are excreted through feces, with studies showing daily excretion rates of around 33 mg in pregnant individuals ^16^. The metabolism of various steroids by gut bacteria not only influences their levels and activity in the body but also transforms them into metabolites with altered bioactivity.

This interaction is particularly evident in the context of sex hormones, where the microbiome’s composition varies by sex and can influence conditions like polycystic ovary syndrome and postmenopausal osteoporosis ^17–20^. Progesterone, a key sex hormone crucial for regulating the menstrual cycle and maintaining pregnancy, is directly synthesized from pregnenolone. The gut microbiome interacts with progesterone in the gut leading to changes in microbial composition ^21–23^, impacting host progesterone levels ^24^, and effecting host reproductive development ^24^. Progesterone levels are tightly regulated and disruptions in the homeostasis of progesterone have been linked to multiple mental and physical disorders ^12^. The potential impact of the microbiome on the levels of progesterone and other steroid hormones could profoundly affect human physical and mental health, especially for pregnant and menstruating individuals.

Progestins, analogs and derivatives of progesterone, are commonly prescribed as therapeutics for postmenopausal symptoms ^10,11^ and as oral contraceptives ^13^. Previous studies have demonstrated that once in the gut, common progestin medicines, including medroxyprogesterone acetate, norethindrone acetate and levonorgestrel can be metabolized by gut microbiome, influencing their efficacy, bioavailability, and safety ^25,26^. A recent study has also demonstrated that progestin metabolites (*e*.*g*., allopregnanolone) can be produced through the microbial metabolism of glucocorticoids in the gut, significantly impacting their concentrations in the guts of pregnant individuals ^27^. The metabolism of progestins by the gut microbiome is an important consideration for their application as therapeutics and for understanding the health of pregnant individuals.

Here, we focus on the reduction of progesterone and related steroid hormones by gut bacteria. *Clostridium innocuum* ^28^ and *Eubacterium* species ^29^ have been found to reduce progesterone into epipregnanolone (3β,5β-Tetrahydroprogesterone), while *Clostridium paraputrificum* can reduce progesterone into pregnanolone (3α,5β-Tetrahydroprogesterone) ^28^. The importance of these biotransformations are highlighted by the differences in functions between progesterone and its derivatives. Progesterone primarily acts through its binding to progesterone receptors, regulating host menstruation and pregnancy ^30^, while epipregnanolone and pregnanolone primarily interact with GABA_A_ receptors, acting as neurosteroids that have impacts on host mood and neuropathology ^31^. Microbial metabolism of progesterone by *C. innocuum*, a medically significant and prevalent gut microbe ^32^, in the gut has been suggested to have a direct impact on serum progesterone levels and follicular development in mice ^24^. Despite the importance of these functions, we still have limited understanding of the microbial genes involved in steroid metabolism in the gut. Crucially, the enzymes responsible for the pathway that reduces progesterone to its 5β reduced and tetrahydroprogesterone derivatives have remained unidentified.

In this study, we characterized multiple microbial enzymes responsible for steroid hormone metabolism in the mammalian gut environment. We first identified a Δ^4^-3-ketosteroid 5β-reductase enzyme family that catalyzes the reduction of progesterone to 5β-dihydroprogesterone, filling a knowledge gap in this pathway in the gut microbiome. We found that this 5β-reductase can act on a variety of steroid hormones, including cortisone and multiple progestins, and identified evolutionary changes in key residues that support its specialized metabolism of steroid hormones. Additionally, we identified a novel 3β-hydroxysteroid dehydrogenase/Δ^5-4^ isomerase capable of converting pregnenolone to progesterone as well as transforming 5β-dihydroprogesterone into epipregnanolone. Interestingly, this gene is often naturally fused with a 5β-reductase, streamlining the entire pathway from pregnenolone to epipregnanolone. We also characterized a Δ^6^-3-ketosteroid reductase enzyme family that reduces 6-dehydroprogesterone to progesterone, which has evolved to specialize in steroid hormones, paralleling the 5β-reductase identified in the study. We performed a metagenomic survey on 1549 samples and observed that 5β-reductase and 3β-hydroxysteroid dehydrogenase/Δ^5-4^ isomerase, while relatively prevalent in the general population, are more prevalent and abundant in females. The enzymes described in this study highlight the diverse mechanisms through which gut bacteria have evolved to metabolize host steroids, providing a molecular basis for understanding how gut microbiota may interact with host hormones and potentially impact menstruation and reproductive health.

## Results

### Comparative genomics uncovers an Old Yellow Enzyme *ci2350* in *Clostridium innocuum* as progesterone 5β-reductase

We identified a putative progesterone 5β-reductase through metabolomic screening and comparative genomics. The proposed progesterone reduction pathway consists of a 5β-reductase enzyme that catalyzes the reduction of progesterone to 5β-dihydroprogesterone, and two additional enzymes, 3α-HSDH (3α-hydroxysteroid dehydrogenase) and 3β-HSDH (3β-hydroxysteroid dehydrogenase) that catalyze the conversion of 5β-dihydroprogesterone to pregnanolone or epipregnanolone respectively (**Figure 1a**) ^28^. Multiple bacterial species from the Clostridia class of the Bacillota phylum have been confirmed to reduce progesterone to 5β-dihydroprogesterone ^28,29^, providing a platform for the identification of putative 5β-reductase enzymes within a collection of Clostridia genomes. We utilized metabolomics to assay for the reduction of progesterone in cultures of potential progesterone reducers and non-reducers. We confirmed that *Clostridium innocuum* 6_1_30 *and Clostridium paraputrificum* NCTC11833 were able to reduce progesterone, corroborating previous findings ^28,29^ (**Figure 1b**).

**Figure 1:**
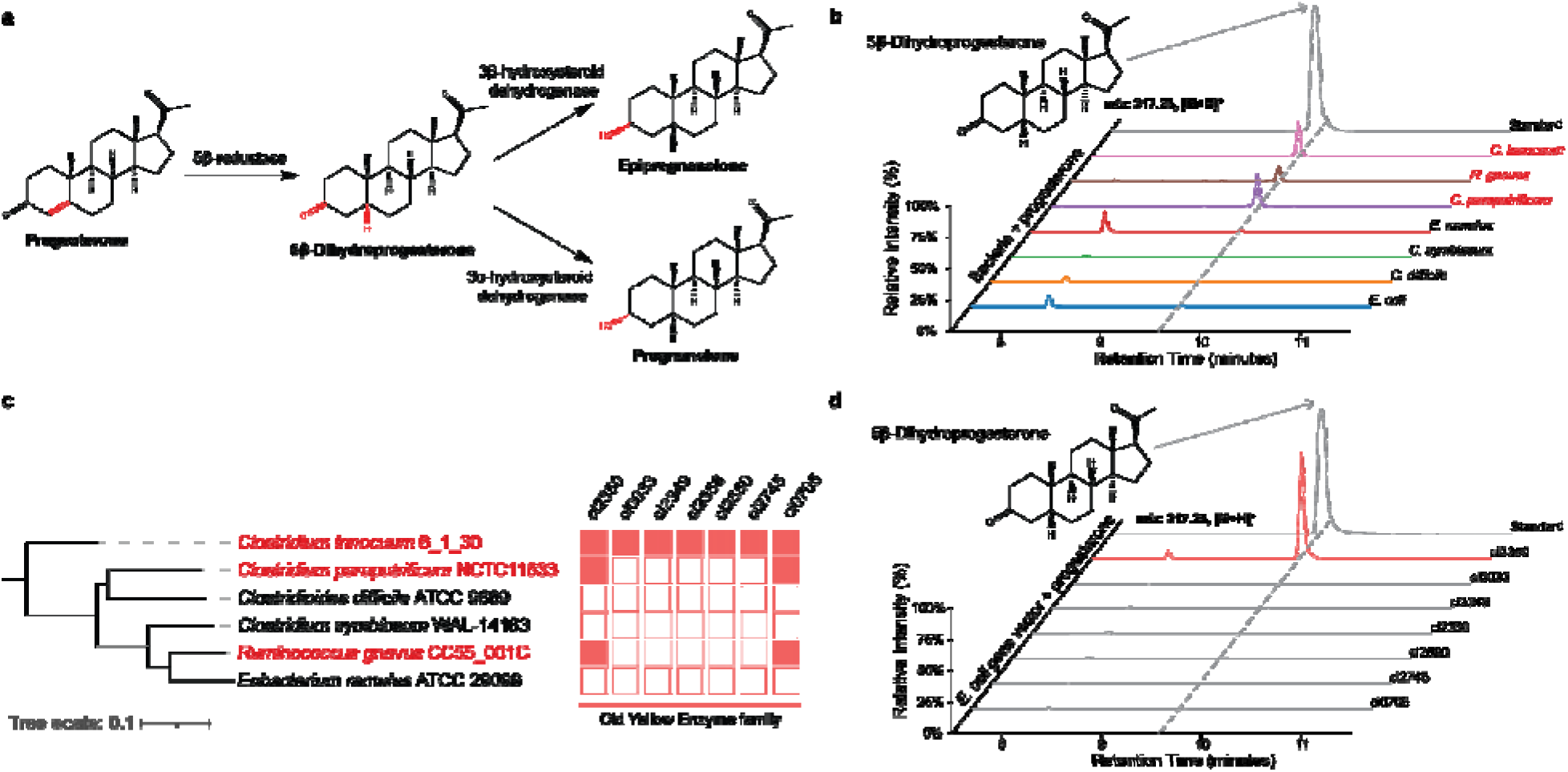
Identification of a progesterone 5β-reductase gene family. **a**, Pathway for the conversion of progesterone to pregnanolone or epipregnanolone. **b**, Detection of 5β-dihydroprogesterone in cultures of seven bacterial strains incubated with progesterone. **c**, Phylogenetic tree and heatmap illustrating the distribution of Old Yellow Enzyme family genes across various bacterial strains. The red-highlighted strains (*Clostridium innocuum* 6_1_30, *Clostridium paraputrificum* NCTC11833, and *Ruminococcus gnavus* CC55_001C) indicate strains with detected enzymatic activity of 5β-reductase. In the heatmap, filled red boxes represent the presence of the corresponding gene, while empty red boxes represent the absence of the corresponding gene. **d**, Comparative chromatograms showing the presence of 5β-dihydroprogesterone in the cultures of the *ci2350* transformed *E. coli* with the strongest activity (highlighted in red) compared to other old yellow enzymes transformed *E. coli*. The chromatograms demonstrate the distinct peaks corresponding to 5β-dihydroprogesterone, confirming the enzymatic activity of *ci2350*.

*Ruminococcus gnavus* (*Mediterraneibacter gnavus)* C55_001C, a common microbe in the human gut, was also able to reduce progesterone demonstrating a previously unknown function of this bacteria that has been associated with various facets of human health and disease ^33,34^ (**Figure 1b**). We did not observe any reduction of progesterone in cultures of *Eubacterium ramulus* ATCC29099, *Clostridium symbiosum* WAL-14163, *Clostridiodes difficile* ATCC9689, or *Escherichia coli* IM93B (**Figure 1b**).

We reasoned that the putative progesterone 5β-reductase genes should be present in the progesterone reducing species and absent in the species that did not reduce progesterone. Previous characterization of Δ^4-3^-ketosteroid 5β-reductases from *C. innocuum* and *C. paraputrificum* has suggested that the enzymes utilize NADH, FMN, and likely FAD as cofactors, and they likely contain iron-sulfur (Fe-S) clusters ^28^. Combined with the fact that the 5β-reductase is proposed to be an ene-reductase reaction (**Figure 1a**), we hypothesized that 5β-reductase would be part of the old yellow enzyme family (COG1902), a family of enzymes that often catalyze stereospecific reduction reactions and are involved in many biologically and medically significant reactions ^35,36^. We identified seven putative old yellow enzymes in the *C. innocuum* genome, two of which had orthologs in *C. paraputrificum* and *R. gnavus* and were absent in the progesterone non-reducing species (**Figure 1c**). To identify if any of these enzymes had 5β-reductase activity, we transformed *E. coli* strains with each of the *C. innocuum* old yellow enzyme orthologs and tested for progesterone reduction (**Figure 1d**). Two of the genes, *ci2350* and *ci0705* matched the expected presence and absence pattern (**Figure 1c**), but only *ci2350* showed any detectable 5β-reductase activity, confirming that *ci2350* encodes a 5β-reductase enzyme capable of reducing progesterone to 5β-dihydroprogesterone.

### *ci2350* is a selective 5β-reductase for steroid hormones

We investigated the activity of the *C. innocuum* progesterone 5β-reductase on various steroids, finding it mainly acts on steroid hormone metabolites. While progesterone is a probable natural substrate for this enzyme, several metabolites exhibiting analogous structural features and bonding patterns may also serve as potential substrates for progesterone 5β-reductase. Using *E. coli* transformed with the ci2350 *C. innocuum* progesterone 5β-reductase gene, we tested for the reduction of progesterone, eleven metabolites which have the same C4 to C5 carbon-carbon double bond on the A ring of the steroid core as progesterone, including three naturally occurring hormones (**Figure 2a**), and eight synthetic progestins (**Figure 2b**). Among the natural steroids, we observed reduction of progesterone and the two progesterone derivatives, cortisone and hydrocortisone (**Figure 2c**). In contrast, we did not observe any reduction of cholest-4-en-3-one, a cholesterol derivative with a similar α,β-unsaturated ketone structure as progesterone (**Figure 2a**). Cholest-4-en-3-one has a much longer hydrocarbon tail and different stereochemistry on the D ring of the steroid core, potentially preventing progesterone 5β-reductase enzyme from reducing this metabolite. A companion study has also demonstrated that these enzymes have broad activity on steroid hormones including corticosterone, prednisolone and testosterone ^37^. This corroborates previous studies where enzyme extracts from *C. innocuum* were able to reduce progesterone and testosterone, but were not able to reduce cholest-4-en-3-one ^28^.

**Figure 2:**
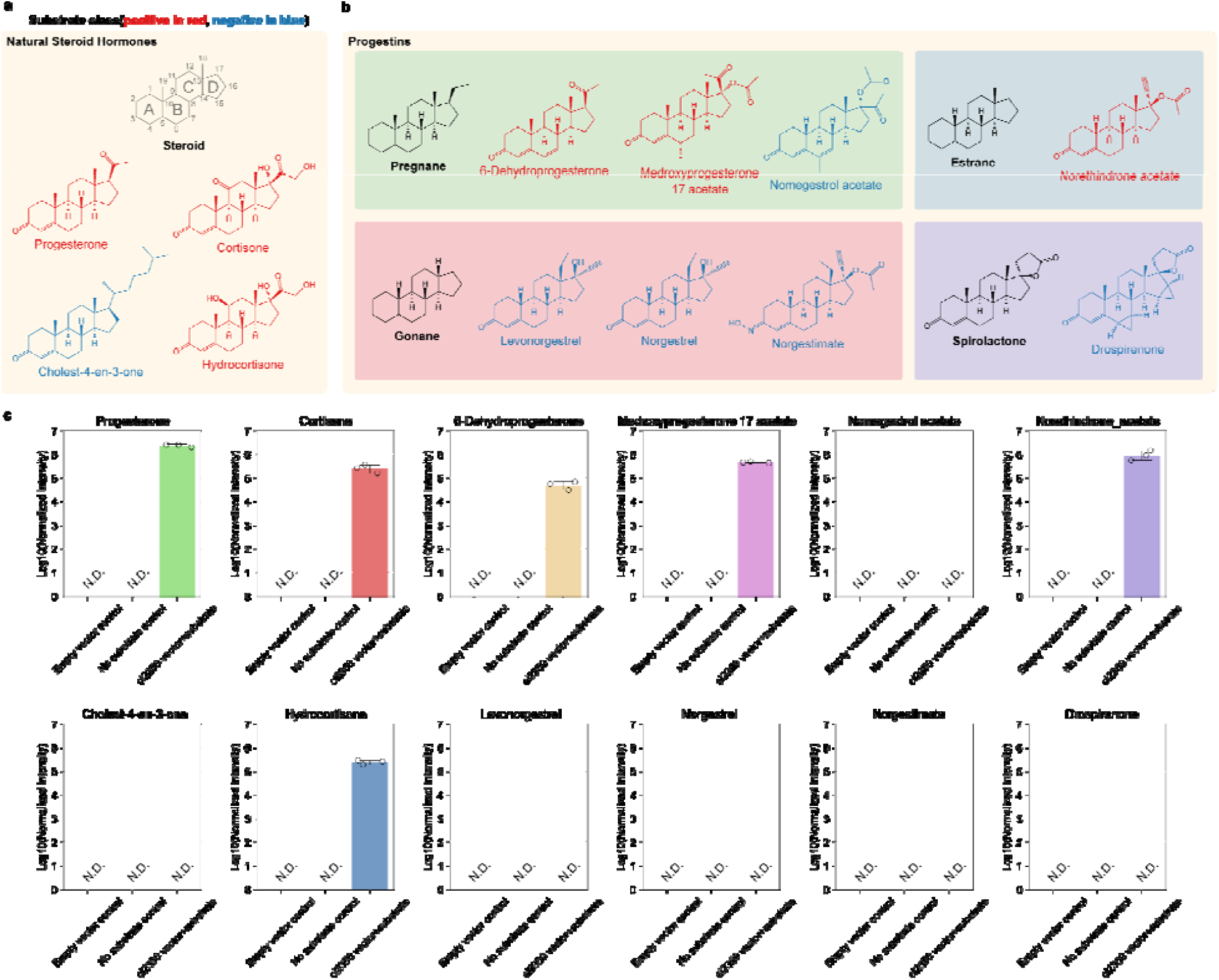
Selective reduction of steroids by progesterone 5β-reductase. **a**, Structures of natural steroid hormones. **b**, Synthetic progestin structures arranged into categories based on their structure and precursors. The steroid core structure with labeled rings is shown in panel a. Backbone structures for different types of metabolites are shown in black in each box. Metabolites colored in red were reduced by ci2350, while metabolites shown in blue were not reduced. **c**, Bar graphs showing mass spectrometry quantified results normalized by intensity from biological triplicates of substrates incubated with cultures of *ci2350* transformed *E. coli*. These are compared to substrates incubated with the *E. coli* transformed with the vector backbone control and a media control.

To test if progesterone 5β-reductase can reduce synthetic progestins we assayed for the reduction of eight progestins derived from four different precursors. Progestin compounds, synthetic progesterone analogs that are often used in birth control and hormonal therapies (Regidor 2014; Prior 2018; Braid and Glasier 1993), can be generally grouped into categories based on what precursors they are synthesized from (*e*.*g*., progesterone, testosterone, 19-norprogesterone) and their activity in the body. When *E. coli* transformed with *ci2350* was incubated with various progestins, we only observed reduction of the progestins in the pregnane and estrane groups, while the gonanes and spironolactone derivatives were not reduced (**Figure 2c**). These differences in reactivity are likely the result of variations in the structure of the metabolites. The metabolites that are reduced share similar structures to progesterone, with methyl groups at the C18 and C19 positions, few modifications to the rest of the steroid ring core, and an unmodified carbonyl group at the C3 carbon (**Figure 2a, Figure 2b**). The exception to this trend is seen in the estrane norethindrone acetate, which is not methylated at the C19 position, but otherwise shares a similar structure to the other reduced metabolites. The remaining metabolites had various differences in structure that could explain the observed lack of reactivity. The gonane progestins lack methyl groups at the C19 position and have an ethyl group instead of a methyl at the C18 position Nomegestrol acetate has an additional α,β, □, □-unsaturated ketone structure and an additional methyl group on the B ring of the steroid core. Norgestimate does not have the typical carbonyl group at the C3 carbon, and drospirenone has two cyclopropanes and a □-lactone modification on the steroid ring structure (**Figure 2b**). These variations in metabolite structure likely contribute to differences in biochemical properties of the metabolites, which may explain the differences in progesterone 5β-reductase activity.

The reduction of specific steroids by progesterone 5β-reductase and lack of activity on others has significant implications for human health. These natural hormones and synthetic progestins have a variety of effects on the human body through their activity as agonists and antagonists of different receptors ^38^. Synthetic progestins are used as therapeutics in fertility and menstrual disorders (*e*.*g*., 6-dehydroprogesterone), as oral contraceptives (*e*.*g*., Medroxyprogesterone 17 acetate, Norethindrone acetate), and as emergency contraceptives (*e*.*g*., levonorgestrel). Notably, Medroxyprogesterone 17 acetate is also effective in treating metastatic breast cancer ^39^. The ability of microbes to metabolize these compounds, altering their bioavailability and producing metabolites with different physiological impacts, has wide ranging implications for the application of these drugs in healthcare settings.

### Natural fusion of 3β-HSDH/Δ^5-4^ isomerase and 5β-reductase mediates conversion of pregnenolone to epipregnanolone

Using the *C. innocuum* 5β-reductase ci2350 protein sequences as a query we identified a putative 5β-reductase clade containing 318 genes from 282 species, primarily from the Bacillota_A and Bacillota phyla (**Figure 3a, Supplementary Table 1**). This includes multiple microbial taxa commonly found in the healthy human gut including members of the Clostridiaceae and Ruminococcaceae family, suggesting that this function may be relatively common in the human gut environment. The 5β-reductase homologs are frequently adjacent to transcription regulators, which vary across the AraC, AcrR, and MerR families (**Figure 3b**). These homologs are also often found next to other enzyme-encoding genes, including baiH homologs and those containing a Rossmann-fold (SSF51735) as a separate gene (**Figure 3b**). Interestingly, the Rossmann-fold (SSF51735) containing gene is fused directly to the 5β-reductase gene at the C-terminus in 61 out of the 257 homologs (**Figure 3b,c**). The Rossmann-fold domain of three genes shares remote homology with the PF01073, the 3β-hydroxysteroid dehydrogenase/Δ^5-4^ isomerase (3β-HSDH/I) of enzymes which is involved in steroid hormone biosynthesis in eukaryotes. In humans the conversion of pregnenolone, the precursor metabolite for steroid hormones ^40^, to progesterone is catalyzed by a 3β-hydroxysteroid dehydrogenase/Δ^5-4^ isomerase ^41^, a function not previously described in bacteria. Based on the similarity to these domains, we suspect that these bacterial 3β-HSDH/I domain containing genes are involved in the metabolism of pregnenolone and potentially 5β-dihydroprogesterone in these bacteria. Given that fused genes are frequently involved in pathway integration ^42^, we hypothesize that the 3β-HSDH/I domain may give these enzymes the ability to catalyze the conversion of pregnenolone to progesterone, 5β-dihydroprogesterone to epipregnanolone, or both in addition to the conversion of progesterone to 5β-dihydroprogesterone that is catalyzed by the 5β-reductase domain (**Figure 3d**).

**Figure 3:**
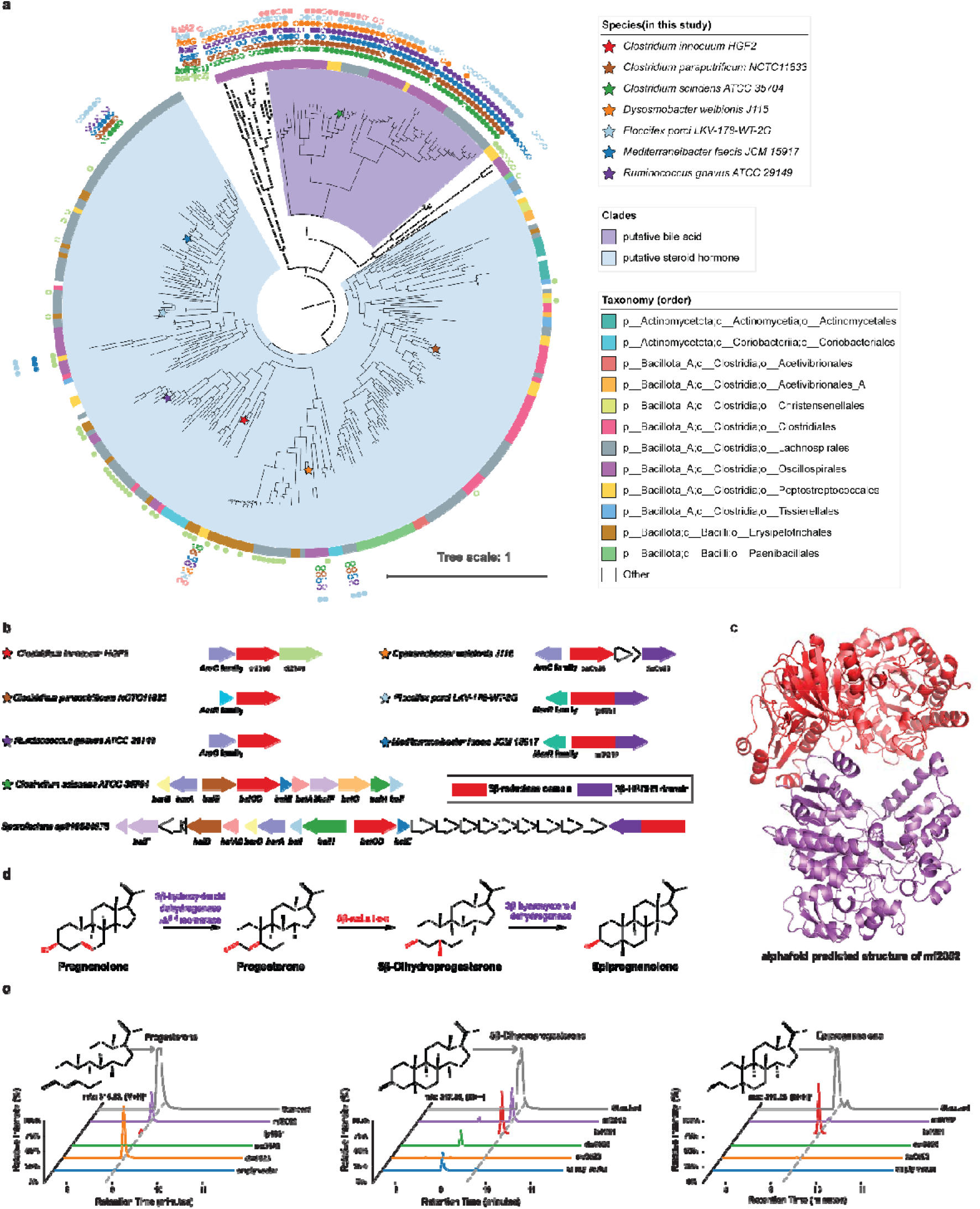
Detection of fused genes composed of 3β-HSDH/Δ^5-4^ isomerase and 5β-reductase; **a**, Phylogenetic tree of putative steroid 5β-reductase clades, rooted by an outgroup. The outer ring indicates the presence of the *bai* operon genes in the same genome: filled circles represent genes in close proximity, while unfilled circles represent genes in the same genome but not in close proximity. The inner ring shows the taxonomic order. Species included in this study are marked with colored stars; **b**, Gene clusters associated with steroid 5β-reductase in various bacterial strains. Arrows represent different genes, with colors corresponding to specific gene functions; **c**. AlphaFold-predicted structure of the fused gene *mf2052* composed of 3β-HSD/Δ^5-4^ isomerase and 5β-reductase, showing the structural domains responsible for each enzymatic activity; **d**, The proposed pathway for the conversion of pregnenolone to epipregnanolone, which could be mediated by different domains of the fused enzyme; **e**, Chromatograms showing the detection of steroid metabolites produced by *E. coli* expressing the genes when using pregnenolone as substrate.

To determine whether the fused genes perform the additional 3β-hydroxysteroid dehydrogenase/Δ^5-4^ isomerase functions, we cloned representative fused genes from *M. faecis* JMC15917 (*mf2052*) and *F. porci* LKV-178-WT-2G (*fp1951*), a 5β-reductase gene from *D. welbionis* J116 (*dw0526*) and separate putative 3β-HSDH/I gene from *D. welbionis J115* (*dw0523*), into *E. coli* and assayed for the conversion of pregnenolone to the potential downstream metabolites progesterone, 5β-dihydroprogesterone, and epipregnanolone. The fused genes (*fp1951* and *mf2052*) and the *D. welbionis* 3β-HSDH/I gene (*dw0523*) were able to metabolize pregnenolone, with *dw0523* producing only progesterone, and the fused genes producing a mixture of progesterone, 5β-dihydroprogesterone, and epipregnanolone (**Figure 3e**). The *D. welbionis* 5β-reductase (*dw0526*) gene alone did not show any activity on pregnanolone, suggesting that pregnanolone conversion to progesterone is catalyzed by the 3β-HSDH/I domain (**Figure 3e**). Overall, this demonstrates that there are multiple routes for pregnenolone metabolism in the gut microbiome, with some bacteria encoding multi-enzyme pathways and others having streamlined pathways catalyzed by fused multi-functional reductases.

We aimed to determine whether the capacity to convert pregnenolone to progesterone is exclusive to the 3β-hydroxysteroid dehydrogenase/Δ^5-4^ isomerase (3β-HSDH/I) family identified in this study or also present in previously recognized 3β-HSDH enzymes. The 3β-HSDH enzyme from *R. gnavus* has been previously shown to perform a 3β-dehydrogenation reaction on bile acids ^43^, which is akin to the conversion of 5β-dihydroprogesterone to epipregnanolone. Close homologs of 3β-HSDH identified in *C. innocuum* 6_1_30 may explain the phenotype of *C. innocuum* converting progesterone to epipregnanolone. To investigate the functions of these genes, we compared the activity of 3β-HSDH genes from *R. gnavus* C55_001C and *C. innocuum* 6_1_30 to the *D. welbionis* 3β-HSHD/I (**Extended Data Figure 1**). The *D. welbionis* 3β-HSDH/I (*dw0523*), the *R. gnavus* 3β-HSDH, and the *C. innocuum* 3β-HSDH genes were all able to convert 5β-dihydroprogesterone to epipregnanolone, demonstrating that these genes have the same 3β-dehydrogenase function, but only the *D. welbionis* 3β-HSHD/I (*dw0523*) was able to convert pregnenolone to progesterone (**Extended Data Figure 1**). This demonstrates that the 3β-hydroxysteroid dehydrogenase/Δ^5-4^ isomerase function is unique, not seen in other bacterial 3β-HSDH enzymes. To our knowledge, this is the first documented instance of a bacterial enzyme exhibiting 3β-HSDH/I function on steroid hormones, unveiling a novel pathway in microbial steroid metabolism.

The identification of pregnenolone metabolizing microbial enzymes has significant implications for human health. Pregnenolone is a key precursor for other steroid hormones and the conversion of it to other products by microbes could directly impact steroid hormone homeostasis. Additionally, altered pregnenolone levels have been implicated in various psychiatric disorders including schizophrenia and bipolar disorder ^44,45^, suggesting that this microbial metabolic process could have wide ranging impacts on human health. Overall, the characterization of these enzymes demonstrates the presence of multiple steroid hormone reduction pathways in these bacteria, with some bacteria being able to catalyze the entire conversion from pregnenolone to epipregnanolone using one fused gene (*M. faecis* JMC15917 and *F. porci* LKV-178-WT-2G), some catalyzing the same complete pathway using two separate enzymes (*D. welbionis* J116), and some bacteria only being able to convert progesterone, but not pregnenolone, to epipregnanolone (*R. gnavus* C55_001C and *C. innocuum* 6_1_30). The streamlining of this pathway through the activity of a fused gene may allow these microbes to take advantage of host-derived pregnenolone and to metabolize hormones more efficiently, potentially giving them a competitive advantage over other species. The diversity in steroid hormone metabolic pathways, and presence of previously unknown routes for hormone metabolism are important considerations for the development of future targeted therapeutics and administration of hormone-based medicine.

### Delineating steroid hormone 5β-reductases

Using a combination of genomic context analysis, ancestral sequence reconstruction and mutagenesis, we identified putative bile acid reductase and steroid hormone reductase clades within a broader family of steroid reductases. We found that the 5β-reductase clade was phylogenetically related to a clade of *baiCD* genes, which encode enzymes that catalyze the reduction of bile acids, another family of steroid metabolites (**Figure 3a**) ^4^. In contrast to the putative 5β-reductase clade, the *baiCD* clade is part of a relatively conserved operon related to bile acid metabolism (**Figure 3a**). Based on the genomic context and phylogenetic relationships, we grouped the genes based on conserved genomic context into a *baiCD* bile acid reductase clade, a progesterone 5β-reductase clade, and a broader steroid 5β-reductase clade that that encompasses both clades (**Figure 3a, Figure 4a**). In fifteen genomes, genes from both the *baiCD* bile acid reductase clade and the progesterone reductases clade are encoded. An example is *Sporofacians sp910584575*, where a steroid 5β-reductase gene and a *bai* operon are located in close proximity, suggesting that there could be functional differences between *baiCD* and the putative progesterone 5β-reductase clade (**Figure 2b**). In total, the differences in genomic context and presence of both genes in some genomes suggests that these two genes may have diverged in function to act on different substrates.

**Figure 4:**
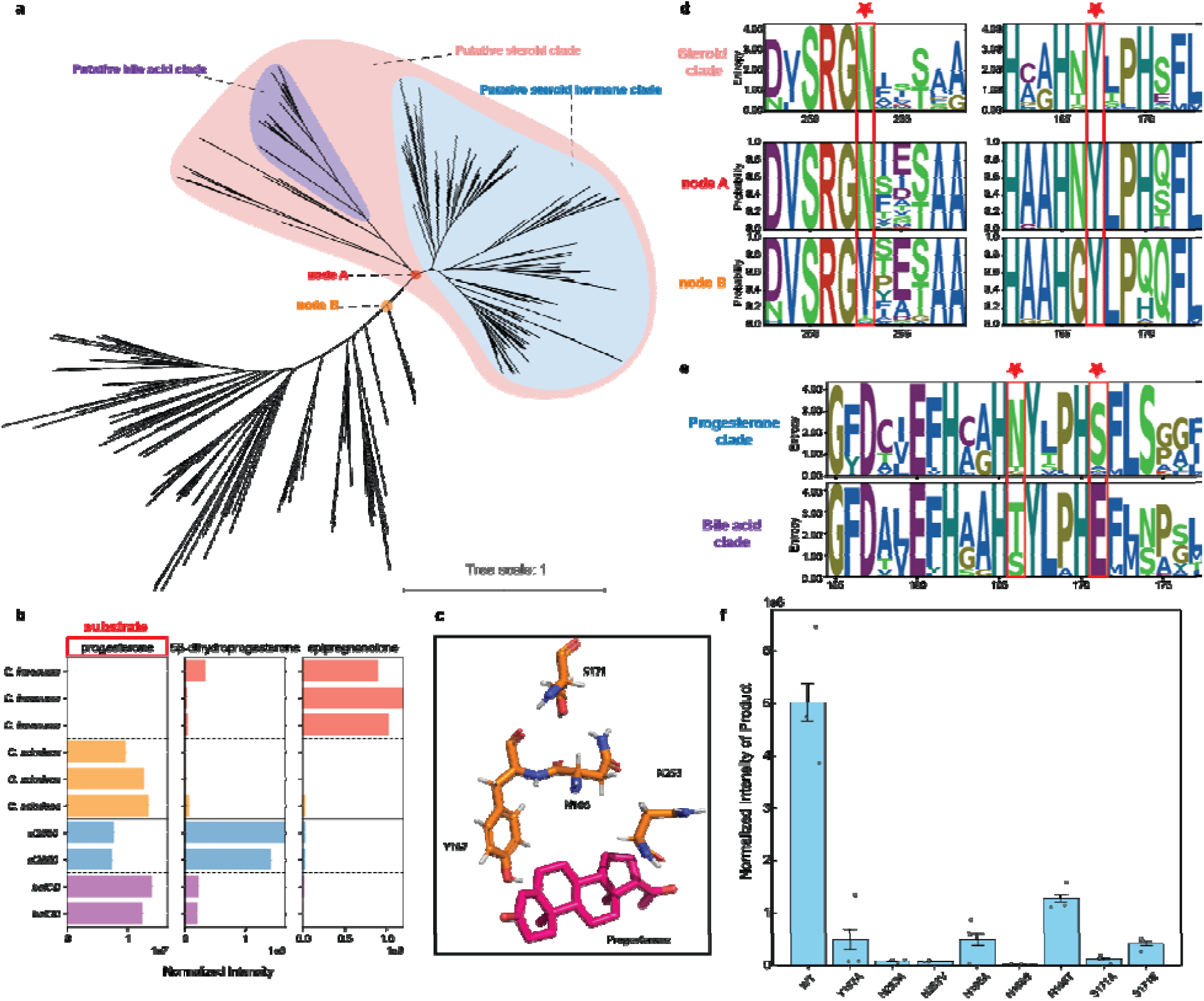
Delineation and subfunctionalization of steroid 5β-reductase. **a**, Phylogenetic tree of Steroid 5β-Reductase related genes, highlighting the putative bile acid clade (purple), putative steroid clade (pink), and putative steroid hormone clade (blue). Nodes A and B are marked, representing key divergence points in the tree. **b**, Bar plots showing the detection of steroid metabolites produced by *Clostridium innocuum HGf2* (red), *Clostridium scindens ATCC 35704* (orange), and *E. coli* transformed with *baiCD* (purple) and *ci2350* (blue). The substrate used is progesterone (highlighted in red box). Each bar panel corresponds to the total ion count of metabolites (progesterone, 5β-dihydroprogesterone, epipregnanolone) detected in LC-MS. **c**, 3D structure of the active site of the enzyme with the substrate progesterone bound, highlighting the key residues involved in substrate binding and catalysis. **d**, Sequence logos for catalytic residues for the Steroid 5β-Reductase clade and the ancestral state reconstructed probability of the putative Steroid 5β-Reductase (nodes A) and the outgroup (node B) (highlighted with red stars). **e**, Sequence logos for key differential residues between the progesterone clade and the bile acid clade (highlighted with red stars). **f**, Bar graph showing the normalized intensity of 5β-dihydroprogesterone detected in the identified enzymes across different mutations, indicating product abundance.

Considering their apparent evolutionary relationship, we next sought to determine if there was functional overlap between the two clades by testing the ability of *C. innocuum* and *C. scindens* to metabolize progesterone. Previous studies have shown that *C. scindens* is capable of metabolizing some bile acids and hormones, but it has been reported to not metabolize progesterone ^46^. Cultures of *C. innocuum* were able to convert progesterone into a mixture of nearly all epipregnenolone with a small amount of detectable 5β-dihydroxyprogesterone remaining (**Figure 4b**). In contrast, the *C. scindens* cultures showed very little activity, with only trace amounts of both potential products being detected (**Figure 4b**). Cloning of the two genes, *C. innocuum ci2350* and *C. scindens baiCD*, into *E. coli* confirmed that these gene products were responsible for the progesterone reductase activity, and showed that *baiCD* had limited, but detectable, reduction of progesterone (**Figure 4b**). While the *C. scindens baiCD* gene may still be capable of catalyzing steroid hormone reduction, differences in cellular transport and regulation may account for the lack of activity seen in the *C. scindens* cultures (**Figure 4b**). Based on these results, we hypothesize that the separate clades within this family of enzymes represent an overall steroid 5β-reductase family that has undergone subfunctionalization to metabolize different substrates.

To delineate the steroid 5β-reductase family from other enzymes, we first aligned the predicted ci2350 to the previously characterized 2,4-dienoyl CoA reductase structure (PDB: 1PS9) and identified plausible regions in ci2350 that could interact with a substrate based on what is known about the active sites of the other enzymes ^47^ (**Figure 4e, Extended Data Figure 2**). Based on this, we identified multiple residues that would have potential interactions with progesterone in the enzyme’s active site (**Figure 4e**). Positions Y167, structurally aligns with a key catalytic residue from the 1PS9 structure (Y166) ^47^, while the N166, S171, and N253 positions are predicted to be in the region of the 5β-reductase enzyme active site (**Extended Data Figure 2**). Through the reconstruction of the predicted ancestral sequences of the steroid 5β-reductase clade, we found that positions Y167 and N253 were conserved in both the steroid hormone and bile acid clades, and N253 was predicted to have undergone a change from valine to aspartic acid in the ancestor of the overall steroid 5β-reductase clade (**Figure 4c, Figure 4d**). We then mutated the ci2350 sequence to change position N253 to a neutral amino acid with no side chain (N253A) and to the predicted ancestral Valine (N253V), finding that N253A lost nearly all of its progesterone reduction activity, while N253V retained a small but detectable level of activity relative to the wild type protein (**Figure 4f**). These results indicate that the steroid 5β-reductase clade can be delineated from other related enzymes at node A in our tree, with the resulting clade representing a functionally distinct family of steroid 5β-reductase enzymes (**Figure 4a**).

With a putative steroid 5β-reductase clade defined, we then identified key differences between the bile acid reductase and steroid hormone reductase clades that contribute to their differences in function. When examining the conservation of the putative active site residues in the steroid 5β-reductase family, we found that the bile acid reductase clade differed from the steroid hormone reductase clade at two potentially interacting positions within the active site, N166, which is primarily tyrosine or serine in the bile acid reductase clade, and S171, which is primarily a glutamic acid in the bile acid reductase clade (**Figure 4d**). Mutants of ci2350 with position N166 mutated to a neutral amino acid with no side chain (N166A) or to serine (N166S), an amino acid with a smaller side chain, showed a nearly complete loss of progesterone reductase activity, while mutation to tyrosine (N166T), the most common amino acid at this position in the bile acid clade, resulted in a reduction of activity to lower, but still detectable levels compared to the wild type (**Figure 4f**). Similarly, mutation of position S171 to an alanine (S171A), a residue with no side chain, or to the most common residue in the bile acid clade (S171E), decreased the progesterone reductase activity of ci2350, to low but detectable levels. The retention of partial activity in mutants reflecting the common residues seen in the bile acid clade highlights the partial redundancy observed when testing the activity of the *C. innocuum* and *C. scindens* proteins. This subfunctionalization within the steroid 5β-reductase family demonstrates an evolutionary pattern where these proteins have evolved to metabolize different but related steroid metabolites.

### A Δ^6^-3-ketosteroid reductase converts a progesterone derivative into its Δ^6^-reduced form

In the *C. innocuum* genome, the 5β-reductase gene (*ci2350*) was adjacent to *ci2349*, which is a homolog of the *baiH*, (**Figure 3b**), the Δ^6^-3-ketosteroid reductases that are known to be involved in bacterial bile acid metabolism ^48^. When performing a broader search for genes similar to ci2349 we identified 94 putative genes. Of the 94 genes identified, 92 are associated with the 5β-reductase gene in the genome, with 78 positioned directly adjacent to it (**Figure 5a, Supplementary Table 2**). These 94 Δ^6^-3-ketosteroid reductase (Δ^6^-RD) genes can be categorized into two clades. The first, Δ^6^-RD(c1), comprises 51 genes that are closely related to the *baiH* gene clade. These genes not only co-localize with the previously identified bile acid reductase clade within the steroid reductase family (**Figure 4a**) but are also predominantly associated with the traditionally defined bai operon ^48,49^. The second clade, Δ^6^-RD(c2), is likely linked with steroid hormone reductases, suggesting specialized functions that may parallel those of the steroid hormone 5β-reductase.

**Figure 5:**
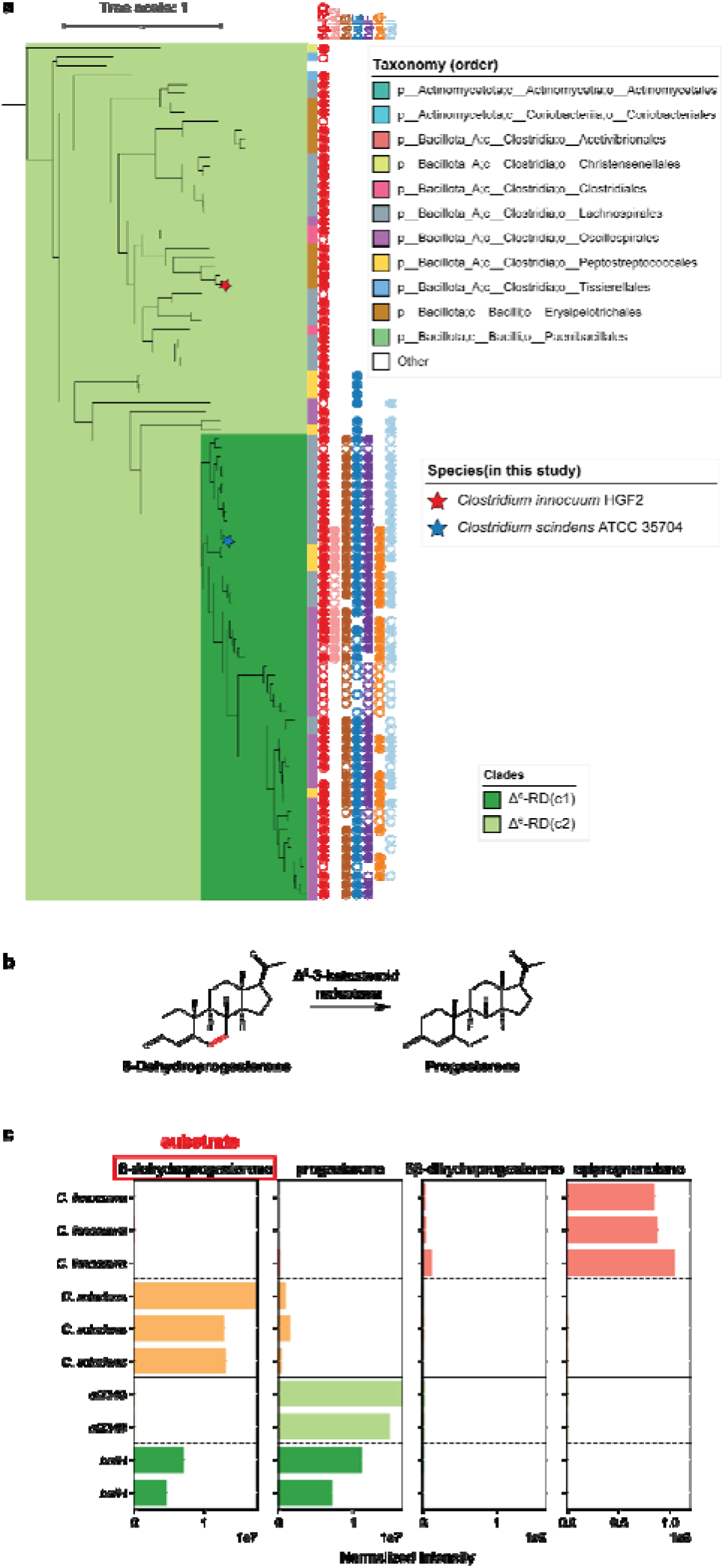
Δ^6^-3-ketosteroid reductase phylogeny and activity in 6-dehydroprogesterone reduction. **a**, Phylogenetic tree of baiH homologs, highlighting clades baiH(c1) and baiH(c2). The circles next to the leaf indicate the presence of the *bai* operon genes in the same genome, with filled circles representing genes in close proximity and unfilled circles representing genes in the same genome but not in close proximity. The bar next to the leaf shows the taxonomic order of the species. Species included in this study are marked with stars: *Clostridium innocuum HGf2* (red star) and *Clostridium scindens* ATCC 35704 (blue star). **b**, Proposed reaction of Δ^6^-3-ketosteroid reductase. **c**, Bar plots showing the detection of steroid metabolites produced by *Clostridium innocuum* HGf2 (orange), *Clostridium scindens* ATCC 35704 (green), and *E. coli* transformed with *baiH* (dark green) and *ci2349* (light green). The substrate used is 6-dehydroprogesterone (highlighted in red). Each bar panel corresponds to the total ion count of metabolites (6-dehydroprogesterone, progesterone, 5β-dihydroprogesterone, epipregnanolone) detected in LC-MS.

To test for differentiation in function between the Δ^6^-RD clades, we assayed for the reduction of 6-dehydroprogesterone, a synthetic progestin, to progesterone. The conversion of 6-dehydroprogesterone to progesterone consists of the reduction of one of the two carbon-carbon double bonds present on the B ring of the steroid nucleus, the same type of reduction catalyzed by BaiH enzymes ^50^ (**Extended Data Figure 3**), suggesting that this is a possible function of the Δ^6^-RD(c2) enzymes (**Figure 5b**). When incubated with 6-dehydroprogesterone, *C. innocuum* showed nearly complete conversion of the 6-dehydroprogesterone to epipregnanolone while *C. scindens* showed only low amounts of conversion to progesterone (**Figure 5c**). When the *baiH* gene from *C. scindens* and the Δ^6^-RD homolog from *C. innocuum* (ci2349) were transformed into *E. coli* we observed efficient conversion of 6-dehydroprogesterone to progesterone by *ci2349*, but only partial conversion of the substrate by the *C. scindens baiH* (**Figure 5c**). This demonstrates that, like the divergence of baiCD and progesterone 5β-reductase within the steroid β-reductase family, this Δ^6^-RD family has undergone subfunctionalization while maintaining partial functional redundancy across the family.

### Differential presence of steroid reductases in male and female gut microbiota

Given the potential role of hormone-metabolizing enzymes in regulating host hormone balance, we then sought to investigate their prevalence and abundance across the human population. Additionally, considering the known differences in steroid hormone production between males and females—which may lead to variations in substrate specificity and availability—and the potential that these enzymes influence health outcomes in a sex-specific manner, we aimed to determine whether their distribution differs between males and females. Therefore, we gathered a total of 1549 metagenomic samples from 13 studies and conducted a comprehensive analysis to assess the presence and abundance of these enzymes across diverse populations (**Supplementary Table 3**).

The presence of Δ^4^-3-ketosteroid 5β-reductase was high, being identified in 94.61% of female samples and 90.59% of male samples, providing evidence that steroid metabolism is a fairly common function in the human gut microbiome (**Figure 6a**). Both 3β-HSDH/I and Δ^6^-RD(c2) were less prevalent but were still fairly common, being found in 82.36% and 57.42% of female and 74.51% and 52.50% of male samples respectively (**Figure 6b,c**). The prevalence of Δ^4^-3-ketosteroid 5β-reductase and 3β-HSDH/I was significantly higher in females compared to males (test of equal proportions; *P=3*.*26e-03, P=2*.*25e-04* respectively) (**Figure 6a,b**), indicating potential differences in the steroid metabolizing microbiome in females compared to males. While the prevalence of Δ6-RD(c2) was not statistically significantly different between sexes, it was more prevalent in female samples, indicating that these trends may be common across different types of steroid metabolism in the gut microbiome. (**Figure 6c**).

**Figure 6:**
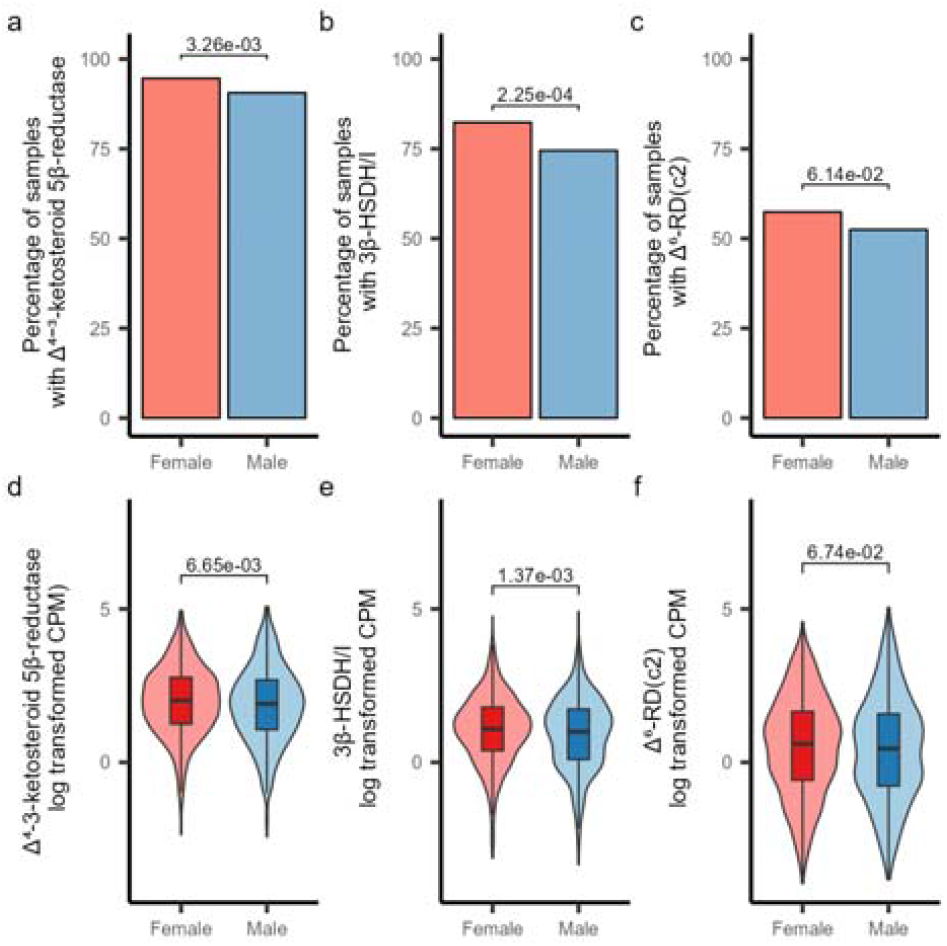
Presence and abundance of steroid hormone reductases in human gut metagenomes. a-c, Plots showing the presence of Δ^4^-3-ketosteroid 5β-reductase (a), 3β-HSDH/I (b), and Δ^6^-RD(c2) (c) in human gut metagenomes. Samples with greater than one CPM mapping to the gene were considered to have the gene present. Statistical comparisons of gene presence are based on a one-sided test of equal proportions to test if the female prevalence was higher than male prevalence. d-f) Plots showing the abundance of Δ^4^-3-ketosteroid 5β-reductase (d), 3β-HSDH/I (e), and Δ^6^-RD(c2) (f) in human gut metagenomes from male and female-derived samples. Plots are shown based on the natural log transformed CPM values (samples with zero count were excluded in the plot). Statistical comparisons are based on one-sided Wilcoxon rank sum tests testing if the CPM values of the female-derived samples are higher than the male-derived samples. All plots show analyses based on a dataset containing 890 female-derived samples and 659 male-derived samples.

In addition to the differences in prevalence, variations in gene abundance were also observed between males and females. The abundances of Δ^4^-3-ketosteroid 5β-reductase and 3β-HSDH/I were significantly higher in females (*Wilcoxon rank sum test; P=6*.*65e-03, P=1*.*37e-03 respectively*), while the abundance of Δ^6^-RD(c2) did not show any difference between sexes (**Figure 6d-f**). While the differences of abundance for Δ^4^-3-ketosteroid 5β-reductase and 3β-HSDH/I were significant, it is unclear if these relatively small differences in abundance would have an impact on the overall steroid metabolizing activity of the gut microbiome. Overall, the differences in the prevalence and abundance of these genes suggests that the sex-related differences in hormone metabolism may influence the presence and abundance of steroid metabolizing bacteria in the human gut.

The observed enrichment of steroid hormone-metabolizing enzymes in females is intriguing, given that these enzymes are also likely to metabolize male hormones such as testosterone derivatives. Even though the exact reasons remain uncertain, several factors could contribute to this pattern: the frequent hormonal fluctuations in females due to menstrual cycles, pregnancy, postpartum and menopause create a dynamic environment that may favor bacteria capable of metabolizing gonadal steroid hormones. Additionally, higher levels of certain hormones, greater exposure to exogenous hormones (e.g., contraceptives), and the need for reproductive health regulation might select for these bacteria. Differences in immune regulation and a more active gut microbiome-hormone axis in females could also play a role in supporting this enrichment. Nevertheless, it highlights the need to investigate how the gut microbiome affects hormone homeostasis, potentially leading to personalized treatments that support both general and reproductive health.

## Discussion

In this study, we identify and characterize multiple novel families of steroid hormone metabolizing enzymes found in mammalian gut bacteria. The identification of these enzymes has highlighted that there are multiple previously uncharacterized routes for bacterial steroid hormone metabolism, involving steroid 5β-reductases, 3β-hydroxysteroid dehydrogenase/Δ^5-4^ isomerase, Δ^6^-3-ketosteroid reductases, and a family of novel multifunction fused genes capable of catalyzing multiple steroid conversions. These enzymes catalyze the biologically significant conversion of multiple steroid hormones and synthetic progestins,exhibiting a relatively high presence in the population and an enrichment in females compared to males. They could potentially impact host physiology, including aspects of female reproductive health, and influence the efficacy of hormone-based medicines such as oral contraceptives, hormone replacement therapy ^31,51,52^. These findings raise new possibilities for understanding the role of the microbiome in human hormone metabolism, particularly in how it affects female reproductive health and hormone-related therapies.

With the identification of these steroid hormone metabolizing enzymes, we can now better understand the impact of the microbiome on hormone homeostasis, reproductive development, menstrual health, pregnancy, menopause, and hormone-related disorders such as polycystic ovary syndrome and endometriosis. Recent reports of the impact of *C. innocuum* progesterone metabolism on follicular development in mice ^24^ and the conversion of corticoids into progestins by bacteria from the Eggerthellaceae family ^27^ highlights the importance of steroid hormone metabolism by the human gut microbiome. Interestingly, a recent study demonstrated that mice that were colonized with pregnancy-associated microbiomes had significantly elevated tetrahydroprogesterone levels, with epigreganolone being the most abundant epimer ^27^. Knowing what microbes and enzymes are involved in the microbial metabolism of endogenous and exogenous steroid hormones has significant and broad implications for advancing our knowledge of female physiology, fertility, hormone-related conditions, and the development of gender-specific medical therapies.

While this study represents an important step forward in understanding the role of the microbiome in steroid hormone metabolism, significant knowledge gaps still exist. Various other factors likely have impacts on the activity and bioavailability of steroid hormones in the gut, including the glyco-conjugation and sulfonation of the metabolites, which have been shown to be impacted by microbial β-glucuronidases ^53^ and sulfatases ^54^. Additionally, the metabolism of progesterone and related metabolites by other groups of bacteria, including Bacteroidetes and Eggerthellaceae, likely impacts the abundance and types of steroids present in the gut. Expanding our knowledge of how these various metabolic processes and bacterial taxa interact will greatly improve our understanding of microbial hormone metabolism and facilitate the translation of this knowledge into medical applications.

Our analysis of the steroid 5β-reductase and Δ^6^-3-ketosteroid reductase enzyme families suggests that bacterial steroid metabolism has coevolved with animal hosts. Steroid metabolism is prevalent across a wide range of life forms, with pregnenolone being produced broadly across vertebrate species ^55^. In contrast, C_24_ bile acids are primarily found in land dwelling vertebrates ^56^, suggesting that their production emerged later in the evolutionary timeline. The evolutionary trajectory of steroids in animal hosts reveals a parallel adaptive shift in the substrate specificity of corresponding bacterial enzymes. It appears that initially broad-spectrum enzyme families involved in steroid metabolism have evolved to include specialized sub-clades, such as baiCD and Δ^6^-RD(c1), which specifically metabolize bile acids. These trends highlight the importance of considering host-context and evolution when trying to understand how the gut microbiome has evolved and the impacts of these microbial enzymes on host physiology.

In conclusion, our study describes the enzymes responsible for steroid hormone reduction in the human gut. These findings highlight the microbiome’s role in human sterol metabolism, which may impact the bioavailability and efficacy of pharmaceutical compounds and alter levels of metabolites being recirculated in the body. The characterization of these enzyme families suggests that bacterial hormone metabolism has evolved in parallel with host metabolism, starting with broad metabolic capabilities and subsequently developing specialized enzymes. Future research will build upon these insights, advancing our understanding of the complex pathways involved in gut microbial biotransformations and their impacts on human health and disease.

## Methods

### Anaerobic bacteria culturing

Bacterial strains were obtained from the NIH Biodefense and Emerging Infections Research Resources Repository (BEI) (**Supplementary Table 4**),including *Clostridium innocuum* 6_1_30, *Clostridium paraputrificum* NCTC11833, *Peptoclostridium difficile*, Strain CD160, *Clostridium symbiosum* WAL-14163, *Ruminococcus gnavus* CC55_001C, *Eubacterium ramulus* ATCC29099, *and Escherichia coli*, K-12, Strain IM93. The strains were inoculated from glycerol stocks and incubated under anaerobic conditions (90% N2, 5% CO2, 5% H2) at 37°C within an anaerobic chamber (Coy Laboratory Products). Growth media consisted of 50 mL of liquid brain-heart infusion (BHI) broth (Research Products International, B11000), supplemented with 1 mg per 100 mL of sterile filtered progesterone or progestins dissolved in ethyl alcohol (EtOH, KOPTEC). Cultures were anaerobically incubated at 37°C for 24-48 hours prior to the introduction of steroid hormones and were then incubated for an additional 48 hours before chloroform extraction.

### UHPLC-MS analysis of steroid hormone

After incubation, the bacterial cultures were centrifuged at 3,260 x g for 10 minutes to pellet the cells. The supernatant was combined with chloroform (4L; Fisher Chemical, 234522) in a separatory funnel for organic extraction. After separation, the chloroform-steroid hormone solution was air-dried under a fan until the chloroform evaporated completely. Subsequently, 1.5 mL of methyl alcohol (4L; Fisher Chemical, 232699). was added to the dried samples and the samples were vortexed to ensure metabolite resuspension. The samples were diluted 1:4 prior to analysis.

Samples were analyzed with the Bruker Maxis-II QTOF, an ultra-high resolution Q-TOF mass spectrometer coupled with a Waters Acquity I-Class PLUS LC system. Liquid chromatography separation was performed on a Phenomenex Kinetex C18 100 Å LC Column (100 × 3 mm, 2.6 μm particle size) with a mobile phase A (water with 0.1% formic acid) and mobile phase B (acetonitrile with 0.1% formic acid). The gradient program was initially set at 10% B in 2 minutes. The linear gradient increased to 90% B in 10 minutes followed by maintenance of 4 minutes before returning to 10% in 1 minute and maintained for 3 minutes. The column was kept at 4°C. The injection volume was 5 μl. Mass spectra were acquired under positive electrospray ionization with an ion spray voltage of 4,500 V. The source temperature (dry gas) was set to 220°C with a flow rate of 5.0 L/min. All data was visualized using the Bruker Compass Analysis software.

### Heterologous expression in E. coli

#### Transformed E. coli Culturing

Transformed *E. coli* strains were cultured by inoculating glycerol stocks derived from agar colony plates into 250 mL of Luria-Bertani (LB) medium (Sigma-Aldrich, 0000324647) supplemented with either 50 μg/mL kanamycin or 100 μg/mL carbenicillin (Chem-Impex, 001453-230228 & Bio Basic, Q3030310). The cultures were aerobically shaken at 37°C for 20 hours. Subsequently, bacterial cells were harvested by centrifugation at 3260 x g for 10 minutes and transferred to an anaerobic chamber. The cell pellet was resuspended in 250 mL of LB medium containing 50 μg/mL kanamycin or 100 μg/mL carbenicillin, along with 1 mg per 100 mL of sterile filtered progesterone, progestins or sterols and 0.4 mM IPTG to induce the expression of the cloned genes. The strains were then anaerobically incubated at 37°C for 48 hours prior to chloroform extraction.

#### Hydrophobic compound preparation

All sterols used in this study were directly dissolved into ethyl alcohol at a concentration of 1 mg per 1 mL and added to media. For LC/MS pregnenolone, progesterone, 5β-dihydroprogesterone, epipregnanolone, 6-dehydroprogesterone, cortisone and hydrocortisone, and all progestins (medroxyprogesterone 17 acetate, levonorgestrel, norgestrel, norgestimate, drospirenone, nomegestrol acetate, norethindrone acetate, ethynodiol diacetate) were suspended in ethyl alcohol and added immediately to BHI or LB broth to undergo reaction. Cholestenone was dissolved in ethyl alcohol and allowed to shake at 220 RPM at 25°C for 18 hours before being added to the media.

#### Transformation with pCW constructs

The *3*β*-HSDH* gene was amplified off of the *C. innocuum* genome, the *3*β*-HSDH* and *3*α*-HSDH* genes were amplified off the *R. gnavus* genome, and the *baiCD* gene was amplified off the *C. scindens* genome using primers designed on Snapgene and ordered on GeneWiz (**Supplementary Table 5**). The amplification was conducted through the polymerase chain reaction (PCR) using OneTaq Master Mix (New England Biolabs (NEB), M0482). The pCW-lic backbone was cut in all instances using a set of two different restriction enzymes, NdeI and HindIII-HF. The amplification product was inserted into the expression vector backbone pCW-lic (Addgene, 26908). A gibson assembly was performed on the restricted backbone and the amplification product (NEB, E2611S).

The constructs cloned into pCW-lic were transformed into NEB 10-beta competent *E. coli* cells following the manufacturer’s protocol (NEB, C3019). The transformed cells were spread onto Luria-Bertani (LB) plates supplemented with 100 μg/mL carbenicillin to select for transformed colonies. To confirm the presence and integrity of the plasmids within the transformed colonies, Oxford nanopore sequencing was conducted by Plasmidsaurus.

The constructs were individually transformed into NEB 10-beta competent *E. coli* cells using the manufacturer’s protocol (NEB, C3019). The cells were plated on Luria-Bertani (LB) plates with 100 μg/mL carbenicillin to select for transformed colonies. The plasmids in transformed colonies were verified through Oxford nanopore sequencing by Plasmidsaurus.

#### Transformation with pET28 constructs

The *ci2350* gene and all its mutants were synthesized using GeneScript’s gene synthesis service (Piscataway, NJ, USA). The DNA sequence was designed with appropriate restriction enzyme sites for subsequent cloning. The sequence was submitted for synthesis to GeneScript and the synthesized DNA construct was subjected to sequence verification using Oxford nanopore sequencing.

The *baiH* gene was amplified off of the *C. scindens* genome using primers designed on SnapGene and ordered through GeneWiz (**Supplementary Table 5**). The amplification was conducted through the polymerase chain reaction (PCR) using OneTaq Master Mix (New England Biolabs (NEB), M0482). The pET28-a(+) backbone was cut using two different restriction enzymes, NdeI and XhoI. The amplification product was inserted into the expression vector backbone (Addgene, 69864-3). A gibson assembly was performed on the restricted backbone and the amplification product (NEB, E2611S).

The synthesized and cloned constructs were individually transformed into NEB T7 Express lysY/Iq Competent *E. coli* (High Efficiency) cells using the manufacturer’s protocol (NEB, C3013I). The cells were plated on Luria-Bertani (LB) plates with 50 μg/mL kanamycin to select for transformed colonies. The plasmids in transformed colonies were verified through Oxford nanopore sequencing by Plasmidsaurus.

#### Transformation with pET11 constructs

The *ci2349* gene was synthesized using GeneScript’s gene synthesis service (Piscataway, NJ, USA). The DNA sequence was designed with appropriate restriction enzyme sites for subsequent cloning, was submitted to GeneScript for synthesis, and the synthesized products were verified using Oxford nanopore sequencing.

The constructs were individually transformed into NEB T7 Express lysY/Iq Competent *E. coli* (High Efficiency) cells using the manufacturer’s protocol (NEB, C3013I). The cells were plated on Luria-Bertani (LB) plates with 100 μg/mL carbenicillin to select for transformed colonies. The plasmids in transformed colonies were verified through Oxford nanopore sequencing by Plasmidsaurus.

#### Identifying Candidate Δ^4^-3-ketosteroid 5β-reductases

Three genomes from the experimentally confirmed progesterone reducers, *C. innocuum* 6_1_30 (GCA_000183585.2), *C. paraputrificum* NCTC11833 (GCA_900447045.1), and *R. gnavus* CC55_001C (GCA_000507805.1), and three genomes from closely related non-progesterone reducers, *C. difficile* ATCC 9689 (GCA_001077535.1), *C. symbiosum* WAL-14163 (GCA_000189595.1), and *E. ramulus* ATCC 29099 (GCA_000469345.1), were downloaded from NCBI. The genomes were annotated with Prokka (v1.14.5). OrthoFinder (v2.5.5) was used to group the predicted protein sequences into orthogroups, using an e-value cutoff of 1e-120. The taxonomic distribution of each orthogroup was profiled to identify orthogroups that were present in the progesterone-reducing species and absent in the non-progesterone-reducing species. Only 13 OGs shared the same taxonomic distribution. Out of those OGs, only 2, ci2350 and ci0705, were annotated as old yellow enzymes, which were investigated further as the putative Δ^4^-3-ketosteroid 5β-reductase.

#### Phylogenetic tree construction

Representative genomes were downloaded from the Genome Taxonomy Database (GTDB, version r214) ^57^ and annotated using Prokka (version 1.14.6) ^58^ to identify protein sequences. Sequences assigned to COG1092 were then identified using eggNOG-Mapper (version 2.1.3) ^59^. Comparative analysis included a BLASTp search (version 2.15.0+) ^60^ using the query sequence ci2350/ci2349 against these identified sequences, setting a limit to the top 500 hits. Sequence alignment was performed using MUSCLE (version 5.1) ^61^. The alignments were subsequently trimmed to retain only regions that mapped to the ‘old yellow enzyme’ domain. Additionally, areas exhibiting more than 95% gaps were removed to enhance alignment quality. Sequences shorter than 500 amino acids for ci2350 homologs and 450 amino acids for ci2349 homologs were excluded using SeqKit (version 2.7.0) ^62^ to ensure robustness in the analysis. Phylogenetic analysis was carried out using IQ-TREE (version 2.1.2) ^63^, employing the LG+I+G4 model to handle rate heterogeneities across sites. The reliability of the phylogenetic trees was evaluated using 1,000 ultrafast bootstrap replicates. Trees were visualized using the Interactive Tree Of Life (iTOL) ^64^.

#### Structural Prediction and identifying Catalytic Residues

The structures for the putative 5β-reductase protein *Clostridium innocuum* 6_1_30 (GCA_000183585.2) were predicted using AlphaFold (version 2.2.0) (Jumper et al, 2021). Binding pockets were predicted using fpocket (v4.0) with default parameters. The pockets were compared to the homologous *E. coli* 2,4-Dienoyl CoA Reductase (PDB: 1PS9) to identify putative substrate binding regions and catalytic residues. The structure for progesterone (PubChem compound identifier: 5994) was docked onto the predicted structure of 5β-reductase using AutoDock Vina (v1.2.5) ^65^. The docking simulation was performed within 15□Å□×□15□Å□×□15□Å cubes centered on the center points of the chosen Fpocket ^66^ substrate binding pocket with exhaustiveness set to 32. Docking results were visualized using PyMOL.The 5β-reductase (ci2350) active site residues were identified using TMalign ^67^ on BaiCD and *E. coli* 2,4-Dienoyl CoA Reductase (PDB:1PS9). Based on the current literature, the ci2350 residues that aligned with the known active site residues in 1PS9 were investigated further.

#### Profiling of steroid hormone reductases in metagenomic samples

HMM profiles were built for the 5β-reductase, Δ^6^-RD(c2), and 3β-HSDH/I enzymes based on the identified clades for each gene. First the protein sequences for each gene were aligned using ClustalO (version 1.2.4) ^68^ and then HMM profiles were constructed for each gene set using *hmmbuild* tool from HMMER (version 3.4) ^69^(profiles for each gene are available as part of the ProkFunFind software package available https://github.com/nlm-irp-jianglab/ProkFunFind). For fused Δ^4^-3-ketosteroid 5β-reductase and 3β-HSDH/I, the regions of the sequence corresponding to each of the fused domains were extracted and separately included in the Δ^4^-3-ketosteroid 5β-reductase or 3β-HSDH/I datasets. The HMMER *hmmsearch* tool (version 3.4) ^69^ was used to search for putative hits to each enzyme in the collection of all genomes in GTDB, with the hits being filtered by E-values of 1×10^−240^, 1×10^−220^, and 1×10^−260^ for the 5β-reductase, 3β-HSDH/I and Δ^6^-RD(c2) searches respectively. The nucleotide sequences for the respective hits were then extracted, with fused Δ^4^-3-ketosteroid 5β-reductase and 3β-HSDH/I genes being split into separate parts, and reads were mapped to them in the subsequent metagenomic analysis.

A collection of 890 female and 659 male-derived gut metagenomic samples from 13 studies were used to analyze the presence of the steroid metabolism genes (**Supplementary Table 3**). Each metagenome was downloaded from SRA and adaptor sequences were trimmed from the reads using Trim-Galore with default settings ^70^. The reads were then mapped to the human reference genome (assembly T2T-CHM13v2.0) and potential human contaminant reads were removed using Samtools (version 1.16.1) ^71^. The quality trimmed reads were aligned to the gene reference sets using Bowtie2 (version 2.5.3) ^72^. For biosamples associated with multiple runs, the mapping counts for each gene were summed across all of the runs for that sample. Only samples with at least 1 million reads combined quality trimmed reads per biosample were analyzed further. The read mapping was then normalized to a counts per million (CPM) value for each gene by dividing the count of mapped reads by the total reads and multiplying the value by one million. If a gene had a CPM of greater than 1 in a sample, then the gene was considered to be present in that sample.

## Author Statements

## Author contributions

B.H and X.J conceptualized and supervised the project. G.A., A.J., K.D., S.L., A.Z., J.T.W., M.G., Y.L., B.H., and X.J., performed the experiments and analyzed the data. G.A. and K.D. wrote the original draft of the paper. G.A, K.D., A.J., B.H., and X.J., reviewed and edited the paper.

## Funding information

A.J., K.D., J.T.W., and X.J. are supported by the Division of Intramural Research of the NIH, National Library of Medicine. B.H is supported by startup funding from the University of Maryland and NIH grant 1R35GM155208-01.

## Conflicts of interest

The authors declare that there are no conflicts of interest.

## Acknowledgments

This work utilized the computational resources of the NIH HPC Biowulf cluster. (http://hpc.nih.gov).

## Data and materials availability

The gene information related to the identified Δ^4-3^-ketosteroid 5β-reductases and Δ^6^-3-ketosteroid reductases are available Supplementary Tables 1 and 2 respectively. The information related to the publicly available metagenomics data analyzed in this study are available in Supplementary Table 3. The information related to the bacterial strains used in this study along with their genome information is available in Supplementary Table 4. The information related to the primers used for cloning gene sequences are provided in Supplementary Table 5. Hidden Markov model profiles for the identification of steroid hormone reductases have also been provided as part of the ProkFunFind search tool (https://github.com/nlm-irp-jianglab/ProkFunFind/tree/master/data/Steroid_reductase).

## Supplementary Materials

**Supplementary Table 1: Identified** Δ^**4**^**-3-ketosteroid 5β-reductases**. Information related to the proteins present in the Δ^4-^3-ketosteroid 5β-reductase family. The predicted protein sequence, protein lengths, predicted substrate specificity, and taxonomy are provided for each sequence.

**Supplementary Table 2: Identified** Δ^**6**^**-3-ketosteroid reductases**. Information related to the proteins present in the Δ^6^-3-ketosteroid reductase family. The predicted protein sequence, protein lengths, predicted substrate specificity, and taxonomy are provided for each sequence.

**Supplementary Table 3: Metagenomic sample information**. Information about the metagenomic data analyzed in this study. The table includes the SRA run, sample ids, and associated study information, along with the read mapping counts, and associated sample biological sex metadata.

**Supplementary Table 4: Strain list**. Information about individual strains used during the cloning process. The list includes strain name, BEI strain designation number, Genome ID, and media it was grown in.

**Supplementary Table 5: Primer sequences used for cloning**. Information regarding primers used for gene amplification.This contains the strain, the gene name, the vector backbone, the forward primer sequence, and the reverse primer sequence. The capitalized bases in the Gibson primers are the regions that will anneal to the target sequence for amplification. The lowercase bases represent the additional bases included in the primers to create the necessary overlap with the vector backbone for the Gibson assembly reaction.

## Notes

### Competing Interest Statement

The authors have declared no competing interest.

