## Extended Data for "Gut Bacteria Encode Reductases that Biotransform Steroid Hormones"

### Extended Data Figures

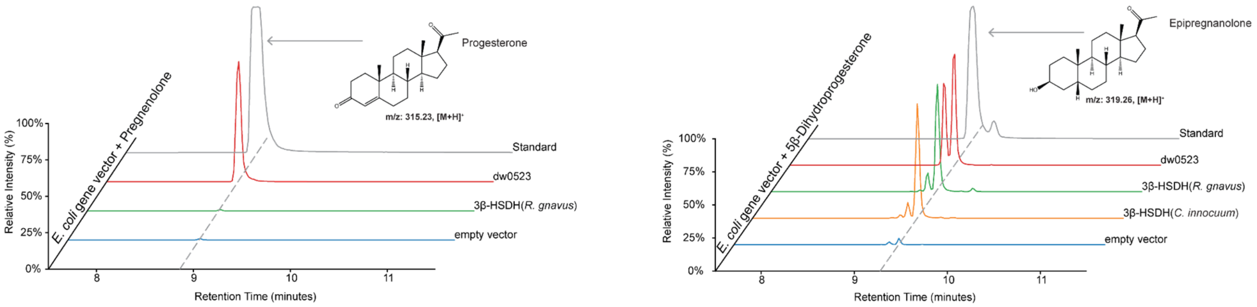


**Extended Data Figure 1: Pregnenolone and 5β-dihydroprogesterone transformation by 3β-HSDH/I and 3β-HSDH transformed *E. coli*.** The left panel shows the ion chromatogram for progesterone (m/z 315.23), while the right panel shows the ion chromatograms for the transformation products 5β-dihydroprogesterone (m/z 317.24) and epipregnanolone (m/z 319.28). Each line color in the chromatograms represents data from *E. coli* transformed with different genes. The structural formulas of the steroids are displayed adjacent to their corresponding peaks, linking the observed chemical transformations with molecular changes.

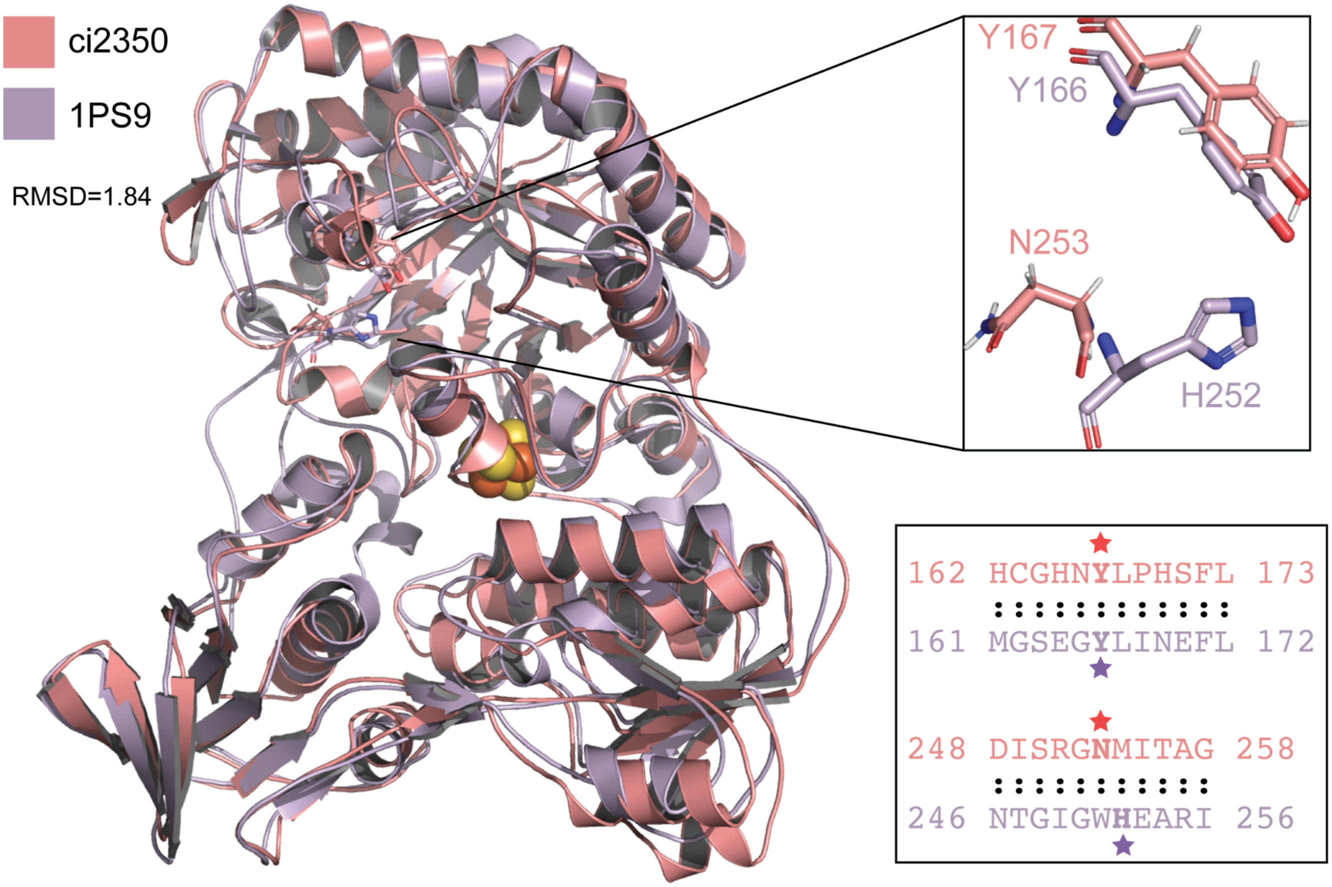


**Extended Data Figure 2: Identification of putative active site residues in ci2350 predicted structure.** Alignment of the AlphaFold predicted structure of the ci2350 steroid hormone 5β-reductase enzyme (red) and the structure of the *E. coli* 2,4-dienoyl CoA reductase enzyme (purple) (PDB: 1PS9). The insert on the top right shows the alignment of the putative ci2350 active site residues with the characterized catalytic residues in the 1PS9 structure. The insert on the bottom right shows subsets of the multiple sequence alignment between the ci2350 and 1PS9 amino acid sequences with stars indicating the positions of the residues of interest in both protein sequences.

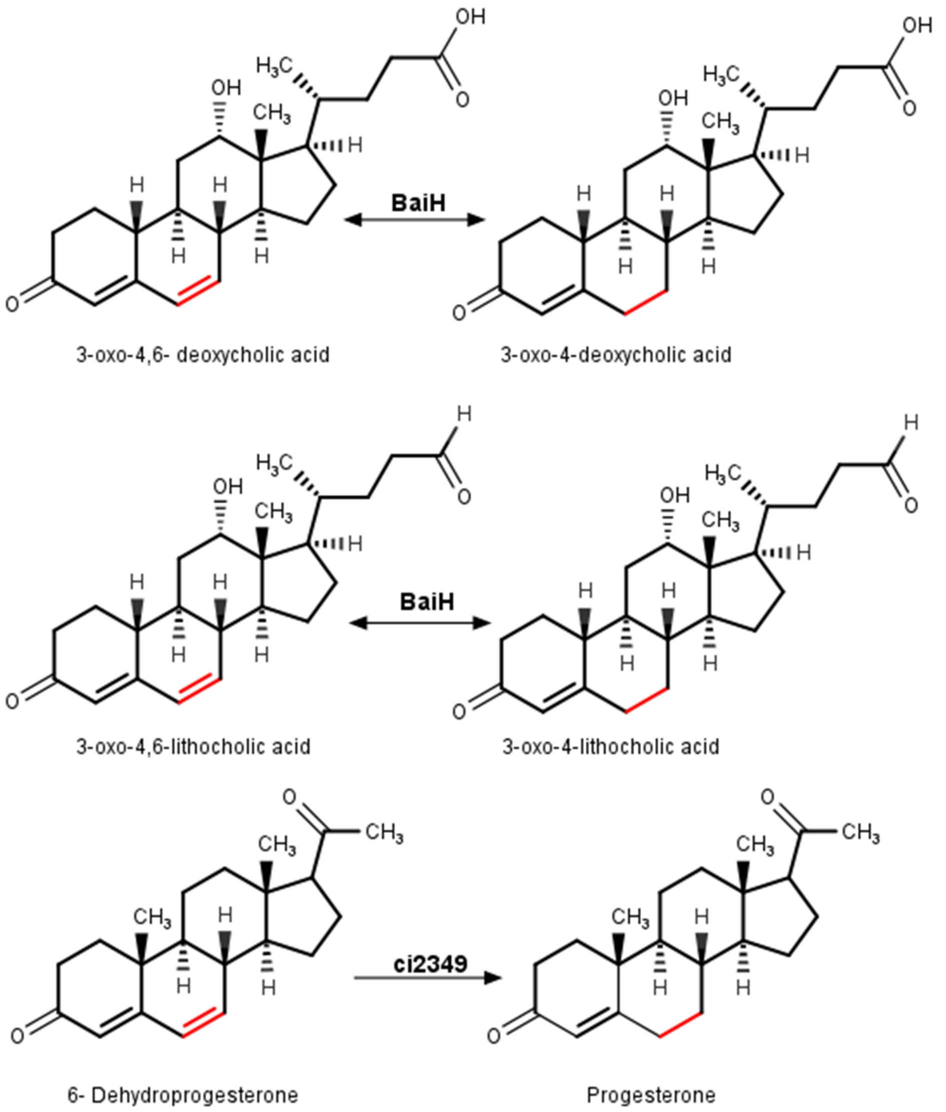


**Extended Data Figure 3: Reactions catalyzed by Δ$^{6}$-3-ketosteroid reductases.** Reaction schematics are shown for the reduction reactions catalyzed by BaiH enzymes (top 2 reactions) and catalyzed by the steroid hormone Δ$^{6}$-3-ketosteroid reductases represented by ci2349 (bottom reaction). The carbon double bonds involved in the reaction are highlighted in red on the left side of the reactions, and the corresponding reduced bond is highlighted in red on the right side of the reactions.
